# Detection of tumor-derived extracellular vesicles interactions with immune cells is dependent on EV-labelling methods

**DOI:** 10.1101/2023.01.04.522609

**Authors:** Luisa Loconte, Davinia Arguedas, Anna Chipont, Rojbin El, Lea Guyonnet, Coralie Guerin, Ester Piovesana, José Luis Vázquez-Ibar, Alain Joliot, Clotilde Théry, Lorena Martín-Jaular

## Abstract

Cell-cell communication within the complex tumor microenvironment is critical to cancer progression. Tumor-derived extracellular vesicles (TD-EVs) are key players in this process. They can interact with immune cells and modulate their activity, either suppressing or activating the immune system. Understanding the interactions between TD-EVs and immune cells is essential for understanding immune modulation by cancer cells. Fluorescent labelling of TD-EVs is a method of choice to study such interaction. This work aims to determine the impact of EV labelling methods on the detection of EV interaction and capture by the different immune cell types within human Peripheral Blood Mononuclear Cells (PBMCs), analyzed by imaging flow cytometry and multicolor spectral flow cytometry. EVs released by the triple-negative breast carcinoma cell line MDA-MB-231 were labeled either with the lipophilic dye MemGlow-488 (MG-488), with Carboxyfluorescein diacetate, succinimidyl ester (CFDA-SE), or through expression of a MyrPalm-superFolder GFP (sfGFP) that incorporates into EVs during their biogenesis using a genetically engineered cell line. Our results showed that these different labeling strategies, although analyzed with the same techniques, led to diverging results. While MG-488-labelled EVs incorporate in all cell types, CFSE-labelled EVs are restricted to a minor subset of cells and sfGFP-labelled EVs are mainly detected in CD14+ monocytes which are the main uptakers of EVs and other particles, regardless of the labeling method. Moreover, MG-488-labeled liposomes behaved similarly to MG-488 EVs, highlighting the predominant role of the labelling strategy on the visualization and analysis of TD-EVs uptake by immune cell types. Consequently, the use of different EV labeling methods has to be considered as they can provide complementary information on various types of EV-cell interaction and EV fate.

## INTRODUCTION

Extracellular Vesicles (EVs) are particles enclosed by a lipid bilayer and secreted by the majority of cells, they play a key role in intercellular communication thanks to their ability to exchange components between cells (1). EVs enclose components such as nucleic acids, lipids and proteins but are highly heterogeneous in composition and size, reflecting the diversity of their biogenesis pathways. Some EVs (exosomes) form first as intraluminal vesicles (ILVs) in multivesicular compartments (generally of endocytic nature), and are secreted upon fusion of these compartments with the plasma membrane. Other EVs are produced directly at the plasma membrane (ectosomes, microvesicles, microparticles, large oncosomes, apoptotic bodies).

EVs play a crucial role in the complex crosstalk within the tumor microenvironment. In particular, tumor-derived EVs (TD-EVs) are able to mediate communication between cancer and immune-cells, promoting either pro-tumoral or anti-tumoral effects. For example, TD-EVs can drive differentiation of monocytes towards myeloid-derived suppressor cells (MDSCs) (2), suppress effector T cell function, generate anergic-like state in natural killer T cells (NKT) (3), promote M2-like macrophage polarization (4), and stimulate regulatory cell expansion (5). Furthermore, NKG2DLs+ EVs can lead to the activation of NK cells, after short time of stimulation, while persistent stimulation leads to sustained NKG2D downmodulation and reduction of NK cell responsiveness (6). Conversely, TD-EVs exhibit immune stimulatory properties under some circumstances. They can transfer major histocompatibility complexes (MHCs) to dendritic cells (DCs), which results in the activation of T cells, thus impairing tumor progression (7). Moreover, TD-EVs bearing macrophage colony-stimulating factor (M-CSF) can induce a unique differentiation signature of monocytes toward pro-inflammatory macrophages, associated with a favorable patient’s clinical conditions (8).

Phenotypic changes triggered by EVs in the recipient cells require uptake of EVs. The mechanism of uptake comprises several sequential steps that are not always concurrent: i) interaction with the recipient cells, ii) entrance in the cells or cell uptake and iii) delivery of EV content to the recipient cell (9). Interaction with and entrance into recipient cells could occur either through specific molecular interactions or through unspecific mechanisms as macropinocytosis (10). EVs can enter into the recipient through endocytosis of the intact vesicle or alternatively, can fuse with the plasma membrane (9). The internalization of EVs has been shown on a wide range of cells such as dendritic cells (11), macrophages (12), dermal fibroblast (13), endothelial and myocardial cells (14). In some cases, cargo delivery has also been demonstrated, but this process could be inefficient depending on the type of recipient cells (15,16).

In order to study TD-EV interaction with immune cells many different EV labelling strategies have been developed. One of the most common method for the study of EV uptake consists in the labelling of EV membrane with fluorescent lipophilic membrane dyes (10,17). Other EV components such as proteins can be targeted using permeable chemical compounds that enter the EV lumen or with the expression of fluorescently-tagged reporters (17). Here, we have compared different approaches of labelling, in order to evaluate the best approach to track EVs capture by recipient cells, monitored by flow cytometry and imaging flow cytometry. Tracking the fate of EVs into immune cells will provide significant insights into immune modulation by TD-EVs in tumor microenvironment and give clues to new therapeutic strategies.

## RESULTS

### The uptake of EVs by PBMCs is time- and temperature-dependent

We used tumor-derived EVs (TD-EVs) from the triple-negative breast carcinoma cell line MDA-MB-231 to test different labelling methods. TD-EVs were obtained from tumor cell conditioned medium (CCM) using size exclusion chromatography (SEC) and collected side-by-side with pools of intermediate and soluble protein fractions as previously described (18). Pooled EVs (fractions 7-11), intermediate (fractions 12-16) and soluble (fractions 17-21) SEC fractions were compared by Western Blot following MISEV2018 guidelines (19). EV fractions from both cell lines contained CD63, CD9, Alix EV markers and were devoid of calreticulin (which was not detected in any fraction), and of 14-3-3, which was present in soluble fractions according to previous publications (8,20). To label EVs with a fluorescent reporter protein (sfGFP-EVs), sfGFP tagged with myristoylation and palmitoylation signals (MyrPalm-sfGFP) was stably expressed in MDA-MB-231 by lentiviral transduction. The resulting protein associates to the inner leaflet of membranes and is incorporated into nascent EVs. We verified that MyrPalm-sfGFP was recovered into EVs secreted by the transduced cells (Figure S1).

EVs secreted by parental MDA-MB-231 were also labeled with two different approaches. First, we labeled the membrane of the EVs with the lipophilic dye MemGlow 488 (MG-EVs) which only fluoresces when inserted into membranes (21). Since this dye is inserted into membranes and could alter the recognition of EV by target cells, we also used CFDA-SE to label MDA-MB-231-derived EVs (CFSE-EVs) (22). CFDA-SE is a membrane-permeable compound that fluoresces after cleavage by esterases present in the lumen of EVs, thus forming CFSE which covalently binds to primary amines inside EVs. As control, fluorescent beads (FB) were used. Fluorescent labelling efficacy of the same number of EVs from the three methods (sfGFP, MG-488, CFSE) or of FBs was quantified using a spectrophotometer (Figure 1A). The same number of TD-EVs or FBs was then incubated with PBMCs for different times (20 minutes, 1 hour and 3 hours) and at two temperatures (37°C and 4°C). The percentage of PBMCs containing fluorescence from EVs or FB (respectively: EV+-PBMCs and FB+-PBMCs) was evaluated by flow cytometry on live cells, using the gating strategy depicted in Figure 1B. Fluorescent EV or FB capture by PBMCs increased with time and was temperature-dependent, since it was reduced at 4°C. However, the percentage of PBMCs containing fluorescence significantly differed between the conditions at both temperatures and most importantly, did not correlate with the intrinsic fluorescence of the particles measured in Figure 1A. FBs, the most fluorescent particles, were modestly uptaken by PBMCs, and their uptake was only partially blocked at 4°C (Figure 1C). Although the labelling intensities of sfGFP-EVs and MG-EVs were similar, but lower than that of CFSE-EVs, the percentage of PBMCs that had incorporated MG-EVs fluorescence was the highest of all groups, while cells having incorporated sfGFP fluorescence were hardly detected (Figure 1C). Surprisingly, the percentage of MG-EV+-PBMCs gradually increased at 4°C, as opposed to the one of CFSE-EV+-PBMCs and sfGFP-EV+-PBMCS which remained very low at this temperature. As most energy-dependent uptake processes are inhibited at 4°C, non EV uptake-mediated MG-488 labeling of target cells might occur. To exclude unspecific labelling of PBMCs by residual free dye present in the EV sample, MG-488 diluted in PBS was processed as MG-EV samples and the same volume was added to PBMCs. No fluorescence was detected in PBMCs incubated with MG-PBS at both temperatures (Figure 1D) ruling out the contamination by free dye after the labelling procedure of EVs. To further investigate the temperature sensitivity of membrane-bound MG-488 dye incorporation, we labelled liposomes with MG-488. Efficient capture by PBMCs was observed with MG-488-labelled liposomes both at 37 and 4 °C, demonstrating that temperature independence was mainly due to the use of MG-488. It suggests that the lipophilic dye MG-488 might diffuse between EVs or liposomes and cells after short/transient contacts.

**Figure 1.**
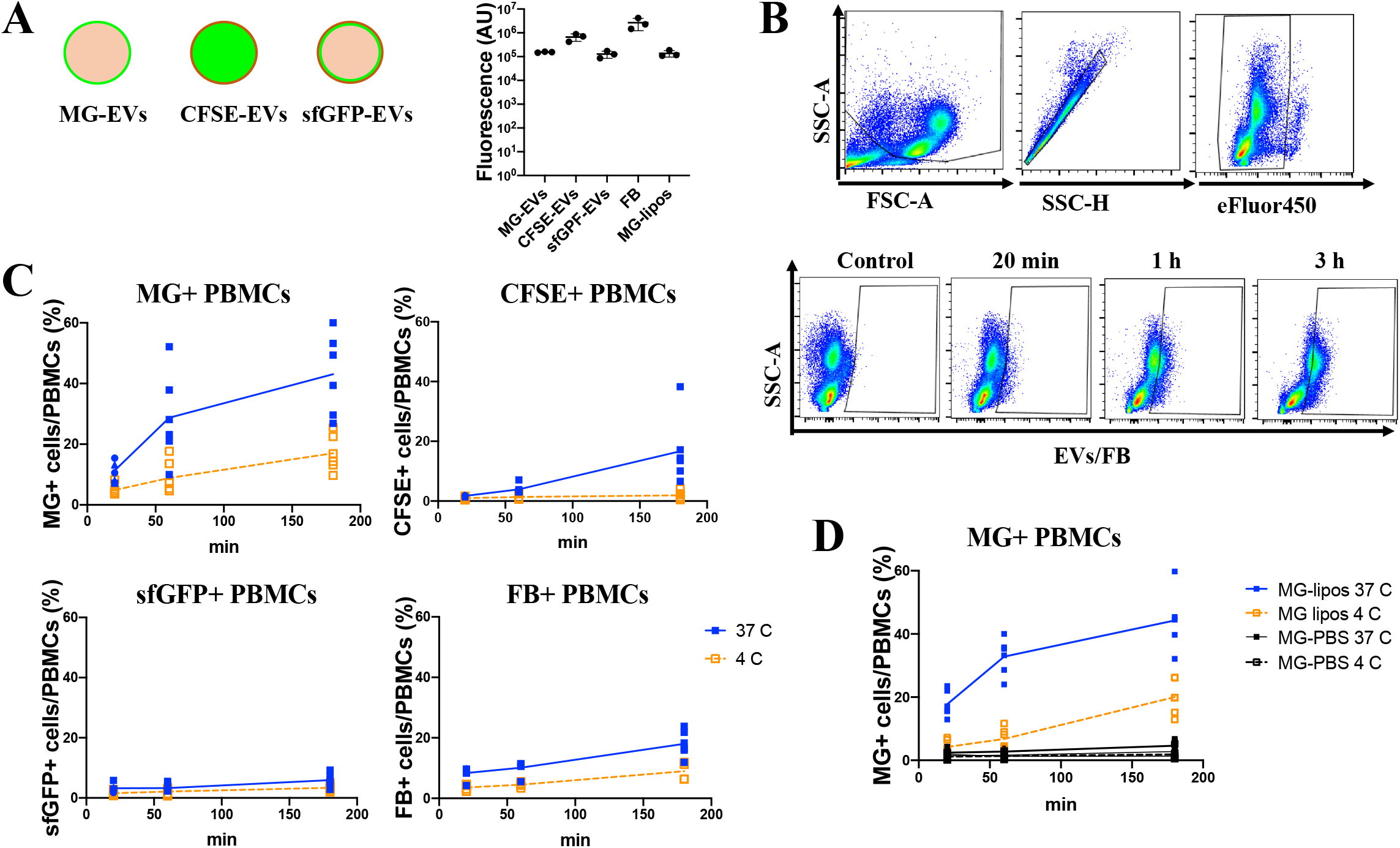
EV uptake by PBMCs is time- and temperature-dependent. (A) The fluorescence of 3×10^8^ MG-EVs, CFSE-EVs, sfGFP-EVs, MG-liposomes or fluorescent beads (FB) was measured with a spectrophotometer (excitation 485/emission 535) prior to their incubation with PBMCs. (B, C) The same number of particles for MG-EVs, CFSE-EVs, sfGFP-EVs or FB was added to 500 000 PBMCs and their uptake by cells was evaluated by analyzing the percentage of MG-488+, CFSE+, sfGFP+ or FB+ PBMCs after 20’, 1hr, 3hrs and either at 37° and 4°C. (B) gating strategy to exclude debris (SSC vs. FSC), then doublets (SSC-A vs. SSC-H) and dead cells (SSC vs. efluor450-viability dye). (C) Each dot corresponds to the uptake by PBMCs from an individual donor. The mean of 4-6 donors is shown. (D) Liposomes labelled with MG-488 and an equivalent volume of PBS incubated with MG-488 and processed in the same way than the labelled EVs were used to assess the uptake by PBMCs at different time points and temperature. The mean of uptake by PBMCs from 6 different donors is shown.

### Detection of the capture and fate of the EVs by PBMCs differs depending on the labelling technique

Next, PBMCs were analyzed by Imaging Flow Cytometry following one-hour incubation with the different fluorescently-labeled particles to analyze their capture and distribution. Cells were defined as focused events that were also singlets, circular and live (Figure S2A). Gating of EV+/beads+-PBMCs was done based on fluorescence (Figure S2B). Consistently to our previous observation (Figure 1C), a large percentage of PBMCs incorporated the MG-EV signal (15 to 100%). The CFSE-EV signal was detected in a smaller but significant percentage of EVs (5 to 60%). By contrast, the percentage of PBMCs that incorporated beads was low (around 5%), while hardly any cells with incorporated sfGFP-EVs were detected (Figure S2B).

We took advantage of the imaging technology combined to flow cytometry to analyze the intracellular distribution of fluorescence in each cell. The texture parameter “Homogeneity” was used to differentiate between cells with dotted (low homogeneity) and diffused (high homogeneity) distributions, to distinguish the retention of fluorescence within intact EV from its release into cells respectively (Figure 2A,2B). We verified that cells incubated with FBs that cannot deliver their content or be degraded were all of low homogeneity values. By contrast, the majority of the cells incubated with MG- and CFSE-EVs presented high homogeneity values, suggestive of a delivery of the EVs content, although dye transfer from EV to the cell membrane in the case of MG could also occur. For both types of labelled EVs, we also detected the presence of cells with low homogeneity values, likely corresponding to intact EVs at the surface of or inside the cells (Figure 2C). By contrast, the small percentage of fluorescent cells detected upon incubation with sfGFP-EVs showed only low homogeneity values. We also noticed that cells presenting low homogeneity values were mainly cells with small area, characteristic of lymphocytes among PBMCs (Figure 2D, left). However, when comparing the percentages of cells that had incorporated EVs/beads (Figure 2D, left) with the frequency of cells with different area within the PBMC samples (Figure S2C), a higher representation in the fluorescent cells of large cells, likely corresponding to myeloid cells, suggested that the latter seemed to preferentially capture all types of particles. This over-representation of cells with large area is even clearer in cells with high homogeneity values that have incorporated CFSE-EV signal (Figure 2D, right), indicating that cells with large area are preferentially capturing EVs. The majority of these CFSE+-cells after 1h were of high homogeneity value, suggestive of the delivery of EV content. In contrast, sfGFP-EVs were not detected in cells with high homogeneity values. Signal dilution, quenching, or rapid degradation of the sfGFP protein within the cell could account for this result. Importantly, cells with high homogeneity values that had incorporated MG-EVs labeling were not very different in size distribution (Figure 2D, right) compared to the total population (Figure S2D), which points to a non-selective transfer of the dye from EVs to the plasma membrane of all PBMCs after short/transient contact with EVs. To address this hypothesis, we distinguished between membrane- and cytosol-associated fluorescent signal in high homogeneity recipient cells incubated with MG-EVs or CFSE-EVs. The membrane/cytosol ratio was close to 1 for CFSE+PBMCS indicating a uniform distribution of EV content in the recipient cell. By contrast, this ratio was greater than 1 in cells that incorporated the MG-EVs signal, indicating that this signal was mostly incorporated at the plasma membrane level.

**Figure 2.**
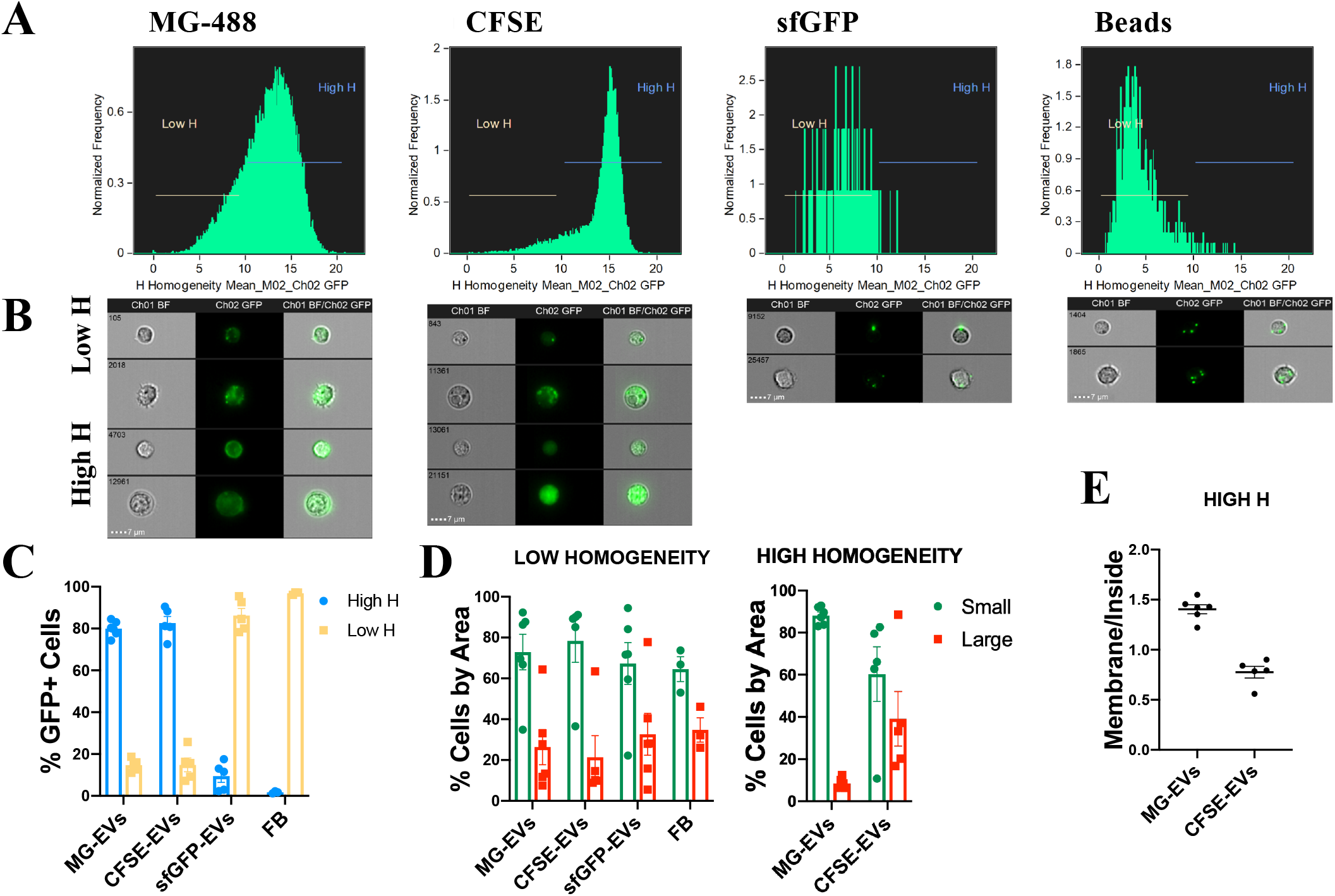
Fate of the different types of labelled EVs in PBMCS after capture. 1 500 000 PBMCs were incubated for 1hr with 9×10^8^ MG-EVs, CFSE-EVs, sfGPF-EVs or FB. (A) Homogeneity mean was evaluated on fluorescent EV+/FB+ cells. (B) Some examples of images of fluorescent EV+/FB+ +cells with high and low homogeneity values are shown. (C) Percentages of cells with high or low Homogeneity values among PBMCs with signal positive for EVs or FB dyes are shown. (D) Low homogeneity and high homogeneity cells were analyzed for their area. (E) Cells with high homogeneity values are analyzed for the location of the signal. Ratio of membrane fluorescence signal to inside signal is shown for PBMCs from 5-6 donors. Each dot corresponds to one PBMC donor.

### CD14+ monocytes are the major cells capturing EVs and beads within PBMCs

Heterogeneous fluorescence distribution between small and large area cells suggested some selectivity of EV cell uptake among PBMC types. To confirm this hypothesis and identify the major immune cells capturing EVs, we used a panel of specific antibody markers to identify major immune subsets within PBMCs using multicolor spectral flow cytometry. After gating single live cells (Figure 1B, upper panel). NK cells were identified as CD3-CD56+, NKT cells as CD3+ CD56+ and T cells as CD3+CD56-. T cells were further defined as CD8+ T cells (CD3+ CD8+) or CD4+ T cells (CD3+ CD4+). B cells were defined as CD3-CD56-CD19+. CD3-CD56-CD19-cells were further classified as classical monocytes (CD14+CD16-), Intermediate monocytes (CD14+ CD16+) or non-conventional monocytes (CD16+ CD14-). Other myeloid cells including cDCs and pDC were defined as CD3-CD56-CD19-CD14-CD16-HLADR+ (Figure S3).

To monitor the capture of TD-EV by each immune subset, identification of cell subtypes was performed first in the whole population of live PBMCs, then, for each identified immune cell type, the EV-associated median fluorescence intensity (MFI) was monitored at different time points of incubation (Figure 3A). For all conditions, the MFI of some immune subsets increased with time up to 3h post-incubation. However, the uptake pattern exhibited by the immune subtypes incubated with TD-EVs significantly differed depending on the EV labeling method. Mainly CD14+CD16-classical monocytes incorporated sfGFP-EVs and CFSE-EVs fluorescence, while the rest of the immune cells were barely labelled for sfGFP-EVs (Figure3A, lower panel). NKs, BCs, intermediate monocytes and other myeloid cells were also labelled after 3h of incubation with CFSE-EVs at 37°C (Figure 3A, middle panel). The uptake of CFSE-EVs by all the cell types analyzed seemed specific as was totally blocked at 4°C (Figure S4). In contrast, when incubated with MG-EVs, the CD14+ cells (including classical and intermediate monocytes) displayed the strongest fluorescence signal, although strong signals were also observed in all the other cells including NKs, NKTs, B cells, non-conventional monocytes and other myeloid cells, with only T lymphocytes displaying hardly detectable EV-derived fluorescence (Figure 3A upper panel). The labelling pattern obtained after PBMC incubation with fluorescent beads was different from that obtained with EVs, since only CD14+ monocytes (classical and intermediate monocytes) were able to capture beads (Figure 3B. lower panel). Surprisingly, the pattern of labelled immune cell populations after MG-EV incubation was similar to the one obtained after incubation with liposomes labelled with MG-488 (Figure 3B, upper panel).

**Figure 3.**
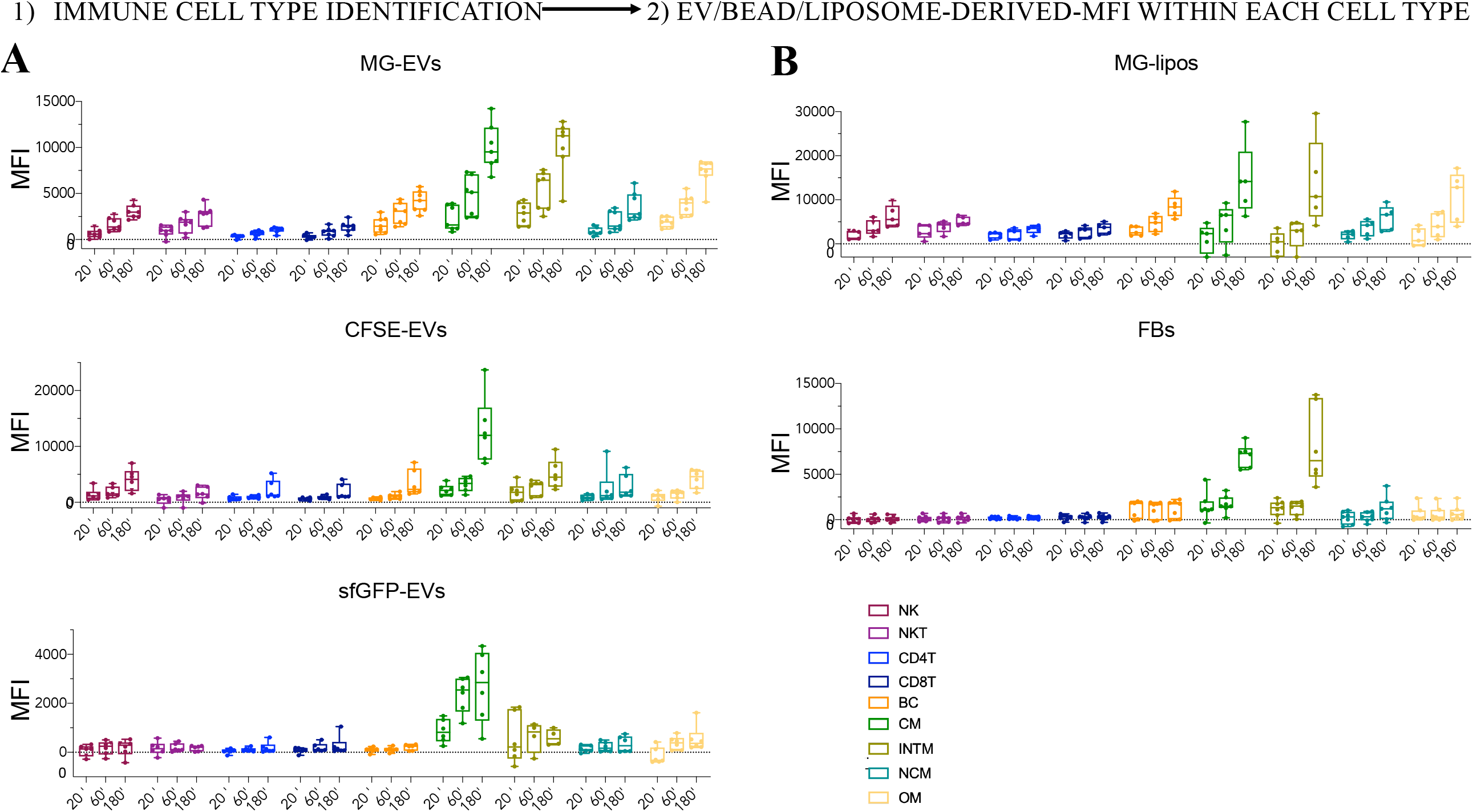
Uptake analysis of EV, beads and liposomes by different immune cells within PBMCs. Differently labelled EVs, beads or liposomes were incubated at 20 min, 1h and 3h at 37°C with PBMCs. Immune cell subtypes were gated into single live cells according to strategy shown in Figure S3. (A) Median Fluorescence intensity (MFI) signal for MG, CFSE or sfGFP was determined in different immune cell types after uptake of TD-EVs and the MFI of a control (no EVs) was subtracted. (B) Median Fluorescence intensity (MFI) signal for FB or MG was determined in different immune cell types after 20 min, 1h and 3h uptake of FBs or MG-liposomes at 37°C. Percentages of different immune subtypes among the EV+PBMCs are shown. Each dot corresponds to PBMCs from different donors. NK = natural killer cells, NKT = natural killer T cells, CD4T = CD4+ T cells, CD8T = CD8+ T cells, BC = B cells, CM; classical monocytes NCM; non-conventional monocytes, INTM; intermediate monocytes, OM: other myeloid.

To evaluate the subsets of PBMCs which preferentially captured labelled EVs or beads after 3 hours of incubation, we first gated the EV+-PBMCs or FB+-PBMCs and then identified the different immune cells subtypes (Figure 4A). We compared the frequency of each immune subtype within the positive cells and within the total PBMCS (Figure 4B, Figure S5). Although T cells were the most abundant immune cells among PBMCs (Figure 4B, Figure S5), classical monocytes (CD14+ CD16-) were the most efficient to capture beads or EVs, independently of the labelling method used (Figure 4A). However, classical monocytes represented around 40% of the cells capturing CFSE-EVs and sfGFP-EVs but only 20% for MG-EVs, close to the percentage of classical monocytes found in the non-labelled PBMCs (Figure 4B). Lymphoid cells, such as CD4+T, CD8+T, B cells and NKT cells were also labelled upon MG-EVs as well as MG-liposomes incubation (Figure 4C). These similarities in immune subtype composition of MG+-PBMCs after incubation with MG-labelled EVs or liposomes, which significantly differ from the ones observed with CFSE/sfGFP EVs, further attests the prevalence of the labeling method in the fluorescence-based analysis of EV uptake.

**Figure 4.**
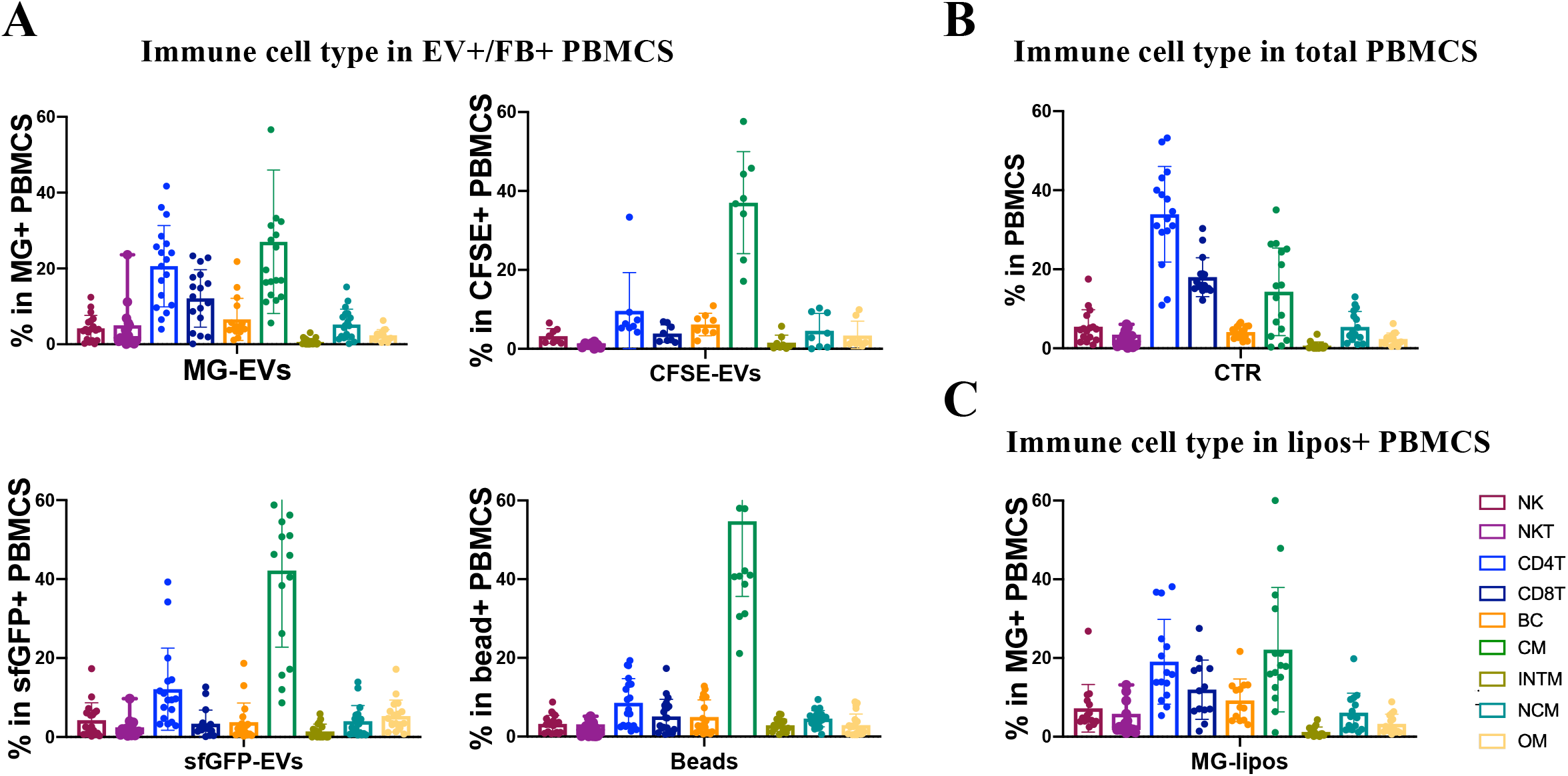
Percentage of immune subtypes among EV+-PBMCs and total PBMCs. Labelled EVs, liposomes or fluorescent beads were incubated for 3h at 37°C with PBMCs and then stained with a panel of antibodies to identify major immune cell populations. MG-488+, CFSE+, sfGFP+ or FB+ -PBMCs were gated to exclude debris, doublets and dead cells (Figure1B). (A) Percentages of different immune subtypes among the EV+/FB+-PBMCs are shown. (B) Percentage of cell subtypes in total PBMCs. (C) PBMCs were incubated for 3h with liposomes and then stained with different antibodies to identify major immune cell populations. Percentages of different subsets within the liposome+-PBMCs are show. Each dot corresponds to PBMCs from different donors. NK = natural killer cells, NKT = natural killer T cells, CD4T = CD4+ T cells, CD8T = CD8+ T cells, BC = B cells, CM; classical monocytes NCM; non-conventional monocytes, INTM; intermediate monocytes, OM: other myeloid.

## DISCUSSION

In this work we have compared three different approaches to label EVs for their capacity to monitor TD-EVs uptake and intracellular fate in PBMCs by spectral flow cytometry and imaging flow cytometry: the lipophilic dye MG-488, the luminal-labelling dye CFSE and the genetically encoded MyrPalm-sfGFP. Regardless of the labelling method used, classical monocytes (CD14+CD16-) are the best uptakers among PBMCS.

Most importantly, our work shows that the detection of TD-EV-associated fluorescence in the recipient cells mainly depends on the labeling method used. Indeed, the three labeling strategies tested in this study are expected to result in different cellular staining patterns after uptake (10). For example, MG-488 is incorporated into the membrane of TD-EVs and, after cell uptake or possibly cell contact, it can be transferred to cell membranes. On the other hand, CFSE and the MyrPalm-sfGFP are inside the TD-EVs that have to be internalized into cells before being able to deliver their content. Importantly, by imaging flow cytometry, we were able to differentiate and quantify cells with likely intact EVs (Low Homogeneity) from those in which the EV content or the lipophilic dye has been transferred (High Homogeneity). Our results show that all labeling strategies allow the detection of intact EVs in a low percentage of EV+-PBMCs. EV content delivery or dye transfer (by membrane contact or fusion) predominates in cells incubated with CFSE-EVs or MG-EVs, respectively. We also observed a strong impact of the EV labelling method on the distribution of fluorescence among the different immune cell types, MG-488 being incorporated in all cell types, CFSE in a minor subset of cells (including lymphoid cells; NK and BC) and sfGFP mainly restricted to CD14+ cells.

The fluorescent properties of MemGlow makes this dye more useful for EV labeling than traditional lipid dyes (21,23). Compared to the other fluorescent dyes used in this study, MG-EV fluorescence is detected in the highest percentage of EV+-PBMCs. However, MG-EV fluorescence is also detected in PBMCs when incubation is performed at 4°C and is not due to free dye contamination. Dye transfer may thus actually occur following simple interaction between the EV and the cell membrane, contrary to CSFE or sfGFP fluorescence. This could lead, at least in part, to a pattern of fluorescence due to redistribution of the dye by normal membrane recycling rather than EV uptake as previously proposed (10,17), which makes interpretations complicated. MG-EV fluorescence is also detected in the majority of cells, including lymphoid cells, NK, NKT, CD8+T and B cells, and the proportion of each cell type among MG-EV+ cells is similar to the one observed in total PBMCs. Importantly, a similar behavior is observed with MG-liposomes, indicating that it does not depend on the specific nature of EVs. In conclusion, MG-488 labeling of EVs does not allow to properly distinguish uptake/content delivery from dye transfer after a brief or transient interaction of the recipient cell with the EVs. Whether or not these transient interactions exclusively detected by MG-488 in our study are relevant to the function of TD-EVs in immune cells remains to be determined.

Our experiments indicate that stable interactions/incorporation of EVs occur mainly in CD14+ cells in accordance with previous publications (8) and consistent with their phagocytic capacity and contribution to particle clearance. However, the uptake of sfGFP+EVs is detected is detected only in a small percentage of PBMCs. Since sfGFP signal is mainly present in cells with low homogeneity values, we hypothesize that we are only able to detect it during the first steps of uptake because after internalization the signal is lost due to dilution, quenching or degradation. Accordingly, our results strongly indicate that CFSE labeling of EVs appears to be the best labeling method to study EV uptake *in vitro*. CFSE allows the detection of intact EVs and their content delivery in a subset of immune cells previously described to incorporate EVs. Importantly, the strict temperature dependence of fluorescence accumulation in PBMCs treated with CSFE-EV clearly supports the requirement for an actual uptake step in these recipient cells. CFSE labelling has been used previously to label EVs (22,24) and does not seem to perturb the size of EVs nor their biodistribution (25). In a recent publication, however, it was described that CFDA-SE was not able to label EVs using protocols inspired by cell labeling (26). In our work we used longer incubation time, in addition to adding a step to remove the free dye after labeling using SEC, as previously described (22).

In conclusion, we have demonstrated in this work that the method used for EV labeling influences the detection of the different types of EV interactions with the recipient cell, including transient EV-PM interaction, EV content delivery and uptake of intact EVs. All these interactions likely occur differently in the various immune cell types and could lead to different functional modifications relevant for communication in the tumor microenvironment.

## ACKNOWLEDGEMENTS

This work was funded by INSERM, CNRS, Institut Curie (including grant from structuring projects call 2021), and grants from European Union Erasmus+ program Student mobility for traineeship 2021/22, and Horizon 2020 research and innovation program under the Marie Skłodowska-Curie grant agreements H2020-MSCA-ITN (722148, TRAIN-EV), and N° 847718 (Institut Curie EuReCa PhD Programme), from french IdEx and LabEx ANR-10-IDEX-0001-02 PSL, french ANR (ANR-18-CE13-0017-03; ANR-18-CE15-0008-01), INCa (INCA_16083), Fondation ARC (ARCPGA12021020003189_3588) and CIC IGR-Curie 1428. This work has received support under the program «Investissements d’Avenir» launched by the French Government. This publication reflects only the author’s view and that the European Research Agency is not responsible for any use that may be made of the information it contains.

## MATERIALS AND METHODS

### Cell culture

MDA-MB-231 cells were verified by short tandem repeat analysis. MDA-MB-231 cells were cultured in Dulbecco’s modified Eagle’s medium (DMEM Glutamax, Gibco), with 10% fetal calf serum (FCS, Gibco) and penicillin-Streptomycin (Gibco). MDA-MB-231 cells expressing Myristoylated-Palmitoylated-superfoldedGFP (MyrPalm-sfGFP) were cultured in DMEM 10% FCS medium under antibiotic selection (2 µg/ml puromycin, ThermoFischer Scientific).

### Plasmids

For the generation of the MyrPalm-sfGFP-encoding plasmid pTCP-MPsfGFP, a synthetic construct encoding sfGFP (accession: ASL68970) was fused in frame at its N-terminus with the acylation sequence of mouse LCK protein (aa 1-10) and at its C-terminus with a P2A-puromycin cassette. The construct was inserted into pTRIP-SFFV at the SrfI-KpnI sites. The SFFV promoter was replaced by the CMV promoter of pCDNA5 vector.

### Lentivirus production and in vitro transduction

For lentivirus production, the packaging cell line HEK293-FT was transfected with pTCP-MPsfGFP together with pSAX2 (12260 Adgene) and pCMV-VSV-G (8454 Adgene) plasmids using the kit *TransIT*®-293 *Transfection* Reagent (Mirus Bio) and following manufacturer’s instructions. After 48 hrs, the viral supernatant was harvested, centrifuged and filtered (0.45 μm filter) and used to obtain MyrPalm-sfGFP-MDA-MB-231 transduced cells. 500 ul of viral supernatant was used to infect 0,2×10^6^ cells on 6 well plate. After 24 hrs, infected cells were cultured in selection media containing puromycin at 2 µg/ml. The expression of GFP was analyzed by microscope and by flow cytometry using Aurora analyzer (Cytek).

### Extracellular Vesicle Isolation

To obtain cell conditioned medium (CCM), 3×10^6^ of MDA-MB-231 or MyrPalm-sfGFP MDA-MB-231 cells were plated per T150 flask with DMEM with 10% FCS. After 48hrs, the cells were washed with PBS and the medium was replaced with DMEM without FCS for 24 hrs. CCM was recovered after 24hrs and centrifuged at 300 *g* for 10 min at 4°C to pellet cells. Cells were counted and viability was measured. After 300 g centrifugation, supernatant was transferred to new tubes and centrifuged at 2,000 g for 20 min at 4°C to discard 2K pellet and then concentrated on a Centricon Plus-70 Centrifugal Filter (Millipore; MWCO 10kDa) by centrifugation at 2,000 g at 4°C until the volume was lower than 500ul. EVs were then isolated by Size Exclusion Chromatography (SEC). Briefly, the concentrated medium was loaded on top of 35 nm qEVoriginal size-exclusion columns (Izon, SP5) and processed according to manufacturer’s protocol. After discarding the void volume (fraction 1-6; 3 mL), 2.5 ml representing the fractions 7 to 11 which contain the EVs were recovered. EVs from MDA-MB-231 and MyrPalm-sfGFP MDA-MB-231 recovered on fractions 7-11, were concentrated using 10KDa cut-off filters (Amicon Ultra-15, Millipore) by centrifuging at 3320 g for 15 min at 4°. The samples were aliquoted and stored at −80ºC before being used in for NTA measurements and uptake experiments.

### Liposome Preparation

Small Unilamellar Vesicles (SUV) composed of Egg yolk L-α-Phosphatidylcholine (EyPC, Avanti Polar), 1-palmitoyl-2-oleoyl-sn-glycero-3-phospho-L-serine (POPS, Avanti Polar) and Cholesterol (Sigma) (50:20:30, mol:mol:mol) were obtained using an extrusion method. Briefly, the appropriate amount of lipids was solubilized in 3 ml chloroform in a round-bottom flask. The solvent was evaporated in a rotavapor in order to form a thin phospholipid film at the glass surface of the flask. Multilamellar Vesicles (MLV) were formed after resuspending this phospholipid film in PBS (pH 8,0) combining vortexing and bath sonication, to reach a final lipid concentration of 20 mg/ml. Finally, SUVs were formed by passing 15 times the MLVs trough a 100 nm-pore polycarbonate membrane (Avanti) using a LiposoFast extruder (Avestin).

### Nanoparticle Tracking Analysis (NTA)

Nanoparticle tracking analysis (NTA) was performed using ZetaView PMX-120 (Particle Metrix) with software version 8.04.02. The instrument settings were 22°C, sensitivity of 85 and shutter of 75. For scatter mode, measurements were done at 11 different positions (five cycles per position) and frame rate of 30 frames per second.

### Western Blot

EV, intermediate and soluble pooled fractions were mixed with Laemmli sample buffer without reducing agent (BioRad). After boiling 5 min at 95°C, samples were loaded on a 4-15% Mini-protean TGX-stain free gels (BioRad). Transfer was performed on Immuno-Blot PVDF membranes (BioRad), with the Trans-blot turbo transfer system (BioRad). Blocking was performed during 30 min with Roche blotting solution in TBS 0,1% Tween. Primary antibodies were incubated overnight at 4°C and secondary antibodies during 1h at room temperature (RT). Development was performed using Clarity western ECL substrate (BioRad) and the Chemidoc Touch imager (BioRad). Membranes were incubated with the following antibodies: mouse anti-human CD63 (1/1000 clone H5C6, BD Bioscience), mouse anti-human CD9 (1/1000 clone MM2/57, Millipore), rabbit anti-human 14–3-3 (1/1000 EPR6380, GeneTex), Alix (1/1000 00013156, Covalab), CR (1/1000 612137, BD Transduction), GFP (1/1000 A11122, Invitrogen). Secondary antibodies were purchased from Jackson Immuno-Research. (HRP-conjugated goat anti-rabbit IgG (H+L), HRP conjugated goat antimouse IgG (H+L) and HRP-conjugated goat anti-rat IgG (H+L), Jackson Immuno-Research)

### MemGlow-488 staining of EVs

EVs isolated by SEC were stained using MemGlow™ 488 Fluorogenic Membrane Probe (Cytoskeleton) used at final concentration of 0.02uM. EVs (1.5-3 ×10^10^ particles/ml) were incubated with the probe for 5’at RT keeping it protected from light. In order to wash the excess of dye from the solution, EVs were diluted with filtered PBS and concentrated using 10Kda filter (Millipore Amicon Ultra 0.5mL Ultracel 10K Centrifugal Filters) at 9000 g, until the volume was reduced to 50ul. NTA was performed after labeling to determine the concentration of particles and their percentage of fluorescence.

### CFSE Staining

EVs isolated by SEC were incubated with 20µM CFDA-SE (Thermofisher) for 2hrs at 37° as previously described (22). To remove the excess of dye, SEC using 35 nm qEVoriginal columns was performed. The CFSE+EVs were collected on fractions 7-10 (2 mL) and the concentration of particles and the % of fluorescence was evaluated by NTA.

### Spectrometer analysis

The fluorescence in each sample was quantified using a spectrophotometer (iD3 SpectraMax microplate reader. Molecular Devices, California, USA). Triplicates containing 3×10^8^ particles from each sample were measured in 96-well flat-bottom black plates.

### PBMCs isolation

Fresh blood pockets from healthy donors were processed the same day of arrival on a Ficoll gradient (Lymphoprep, Greiner Bio-One) to obtain Peripheral Blood Mononuclear Cells (PBMCs). Briefly, blood was extracted from each pocket and poured in filtered falcon tubes containing 15ml of Lymphoprep that have been centrifuged previously (1000g,1 min, RT). The tubes were filled up to 50 ml with PBS and then centrifuged at 800 g without brake during 15 min at RT. The PBMC layer interphase was transferred to new falcon tube containing 50mL PBS. Three washes with PBS were done by centrifuging at 200 g for 10 min at RT. Washed PBMCs were stored overnight in complete RPMI media at 4ºC.

### Uptake Experiments by spectral Flow Cytometry

EVs stained with MG-488 or CFSE, EVs containing sfGFP, fluorescent beads (FluoSpheres 0.1 um yellow-green, Invitrogen, F8803) or liposomes stained with MG-488 were diluted to a concentration of 1.5×10^10^ particles/ml. Quantification of the fluorescence of the different samples was done before the uptake experiment by spectrometer. 500 000 PBMCs were seeded in a 96-well plate in 150 μl of RPMI without FBS and treated with 3×10^8^ particles added at different time points (3hrs, 1hrs, 20’). The incubation was performed both at 37°C or 4°C. The samples were then washed with cold PBS in order to stop the uptake and centrifuged (600g, 5’). Cells were then incubated for 30’ at 4°C with 50 μl of Live Dead (eFluor450, Invitrogen, 65-0863-14, dilution 1/100). After the incubation, cells were washed with cold PBS by centrifuging at 600g for 5’. The cells were incubated for 10 min with 25 µl of FcR Blocking Reagent (Miltenyi Biotec 130-059-901) (pre-diluted 1:50 with FACS buffer (PBS, 0.5% BSA, 2mM EDTA). A mix of antibodies was added in a final volume of 25 µl and samples were incubated for 25’. After antibody staining, samples were resuspended in FACS buffer and acquired using the Spectral Flow Cytometer AURORA Cytek (Cytek Biosciences). After deconvolution, data analysis was performed using the FlowJo program (v10.6). FMOs were used during the optimization of the staining. Monocolors and unstained samples were performed to calculate compensation matrix and establish the gatings.

The antibodies used were: anti-**CD3**, Alexa-Fluor 647 (BD Pharmigen, clone UCHT1, diluted 1/100); anti-**CD4**, Pecy5-A (Biolegend, clone OKT4,, diluted 1/100) ; anti-**CD8**, PE, (Biolegend, clone SK1,, diluted 1/100); anti-**CD11c**, BUV395 (BD Horizon, clone B-LY6, diluted 1/100); anti-**CD14**, Pecy-7 (Biolegend, clone HCD14, diluted 1/100); anti-**CD16**, BUV737, (BD, clone 3G8,, diluted 1/100); anti-**CD19** Alexa-Fluor 700 (BD Pharmingen, clone HIB19, diluted 1/50; anti-**HLA-DR** Alexa Fluor 780 (Invitrogen, clone LN3, diluted 1/200); anti-**CD56**, BUV605 (BD, clone B159, diluted 1/100). All the working concentration of the antibodies were determined after titration.

### Uptake experiments by Imaging Flow Cytometry

1 500 000 PBMCs were incubated for 1hr with 9×10^8^ labelled EVs, or fluorescent beads and then were washed with cold FACS buffer and collected in 1.5 ml Eppendorf by centrifugation at 500g for 6-7’. Adequate PBS and unstained controls were performed in parallel. The pellet was resuspendend in 100 μl of FACS buffer. To exclude dead cells, DAPI was incorporated extemporaneously at 1µg/mL. Samples were analyzed at the single cell level by imaging flow cytometry (ImageStream X MKII, Amnis/Luminex). Acquisition template was set for detecting labelled EVs on channel Ch02 with 488nm laser (100mW), DAPI on channel Ch07 with 405nm laser (50mW) and Ch06 SSC with 785nm laser (30mW). Ch01 and Ch09 were used for Brightfields. Monocolors and unstained were performed to calculate compensation matrix.

Analysis was done with IDEAS analysis software (v6.2). We first selected events in “Focus” (Ch01 Gradient RMS) and then “Singlet” cells (Ch01 Area and Aspect Ratio Intensity). Dead cells and non-circular cells (“Live Circular”) were excluded using respectively DAPI and Circularity feature on brightfield. “Ch02+” gate represent events with presence of labelled EVs. To select uptaking cells, Delta Centroid XY feature for Ch01 and Ch02 assess the distance between center of the cell and EVs signal. “Intern GFP” gate correspond to cells with EVs uptaken. H Homegeneity feature on Ch02 enable the distinction between High (“High H) and Low (“Low H”) Homogeneity of EVs signal. Cells were then selected depending on size and granularity with Ch01 Area and Intensity of SSC (“Small” and “Large”). To obtain the Ch02 Intensity Ratio Membrane/Cytoplasm, a Cytoplasm Mask was created by eroding the M01 Mask of 7 pixels (Erode(M01, 7)) and a membrane mask by dilating the M01 Mask of 1 pixel (Dilate(M01, 1)) and substraction of Cytoplasm Mask for a final defined as (Dilate(M01, 1) And Not Cytoplasm Mask). Those two masks were applied to an Intensity Feature, and Ratio was generated. Gating strategy was also used to evaluated “Small” and “Large” followed by “Ch02+” events plus “Intern GFP” and then “Low H” and “High H”.

