## Supplementary figures for "Detection of tumor-derived extracellular vesicles interactions with immune cells is dependent on EV-labelling methods"

***Figure S1. Analysis of EVs isolated from MDA-MB-231 and MyrPalm-sfGFP MDA-MB-231 cells***. EVs were isolated from CCM of MDA-MB-231 and MyrPalm-sfGFP MDA-MB-231 cells and separated from soluble factors by SEC. (A) Pooled EVs (EVs, 7-11), intermediate (Int, 12-16) and soluble (Sol, 17-21) fractions were collected and analyzed by WB. EV fractions from 20 × $10^6$ cells and pooled intermediate and soluble fractions obtained from 8 x $10^6$ cells were loaded and analysed for the indicated proteins (CR = calreticulin). (*B*) Particle quantification in pooled EV fractions is shown. Each dot represents an independent isolation.

***Figure S2. Gating strategy for ImageStream capture assay****. 1 500 000 PBMCs were incubated for 1hr with* $9x10^8$ *EVs labelled with the three different strategies, or with fluorescent beads. Cells were washed with cold FACS buffer. The samples were analyzed then by imaging flow cytometry (ImageStream X MKII, Amnis/Luminex). (A) Gating strategy is shown in a control sample not incubated with fluorescent EVs or beads. (B) Percentage of EV+ /bead+ cells (= Ch02+) was determined in 4-6 experiments using the gate in (A) and discarding EV signals not coincident with cells using the* delta Centroid XY feature for Ch01 to assess the distance between center of the cell and EVs signal. *(C) Gating of cells according to their size was done on focused live circular singlets. Quantification of small and large cells for the donors used in the different experiments is shown.*

***Figure S3. Gating strategy adopted in order to analyze the different immune subsets within PBMCs.*** *Different immune subsets were identified based on negative and positive expression of markers; NK cells (CD3- CD56+), NKT cells (CD3+ CD56+), CD8+ T cells (CD3+ CD8+), CD4+ T cells (CD3+ CD4+), B cells (CD3- CD56- CD19+), Classical monocytes (CD3- CD56- CD19- CD14+ CD16-), Intermediate monocytes (CD3- CD56- CD19- CD14+ CD16+), Non-conventional monocytes (CD3- CD56- CD19- CD14- CD16+) and other myeloid cells (CD3- CD56- CD19- CD14- CD16- HLADR+), including dendritic cells (HLADR+ CD11c+) and pDCs (HLADR+ CD11c-).*

***Figure S5. Percentage of immune subtypes among total PBMCs.*** *Labelled EVs, liposomes or fluorescent beads were incubated for 3h at 37°C with PBMCs and then stained with a panel of antibodies to identify major immune cell populations. Percentage* *of cell subtypes in total PBMCs. Each dot corresponds to PBMCs from different donors. NK = natural killer cells, NKT = natural killer T cells, CD4T = CD4+ T cells, CD8T = CD8+ T cells, BC = B cells, CM; classical monocytes NCM; non-conventional monocytes, INTM; intermediate monocytes, OM: other myeloid.*

***Figure S4. Uptake analysis of CFSE-EVs by PBMCs at 4$^0$C.*** *CFSE EVs were incubated at different time points at 4°C with PBMCs. Immune cell subtypes were gated into single live cells according to Figure S3. MFI signal for a control samples (no EVs) was subtracted to the CFSE MFI. Each dot corresponds to PBMCs from different donors. NK = natural killer cells, NKT = natural killer T cells, CD4T = CD4+ T cells, CD8T = CD8+ T cells, BC = B cells, CM; classical monocytes NCM; non-conventional monocytes, INTM; intermediate monocytes, OM: other myeloid.*

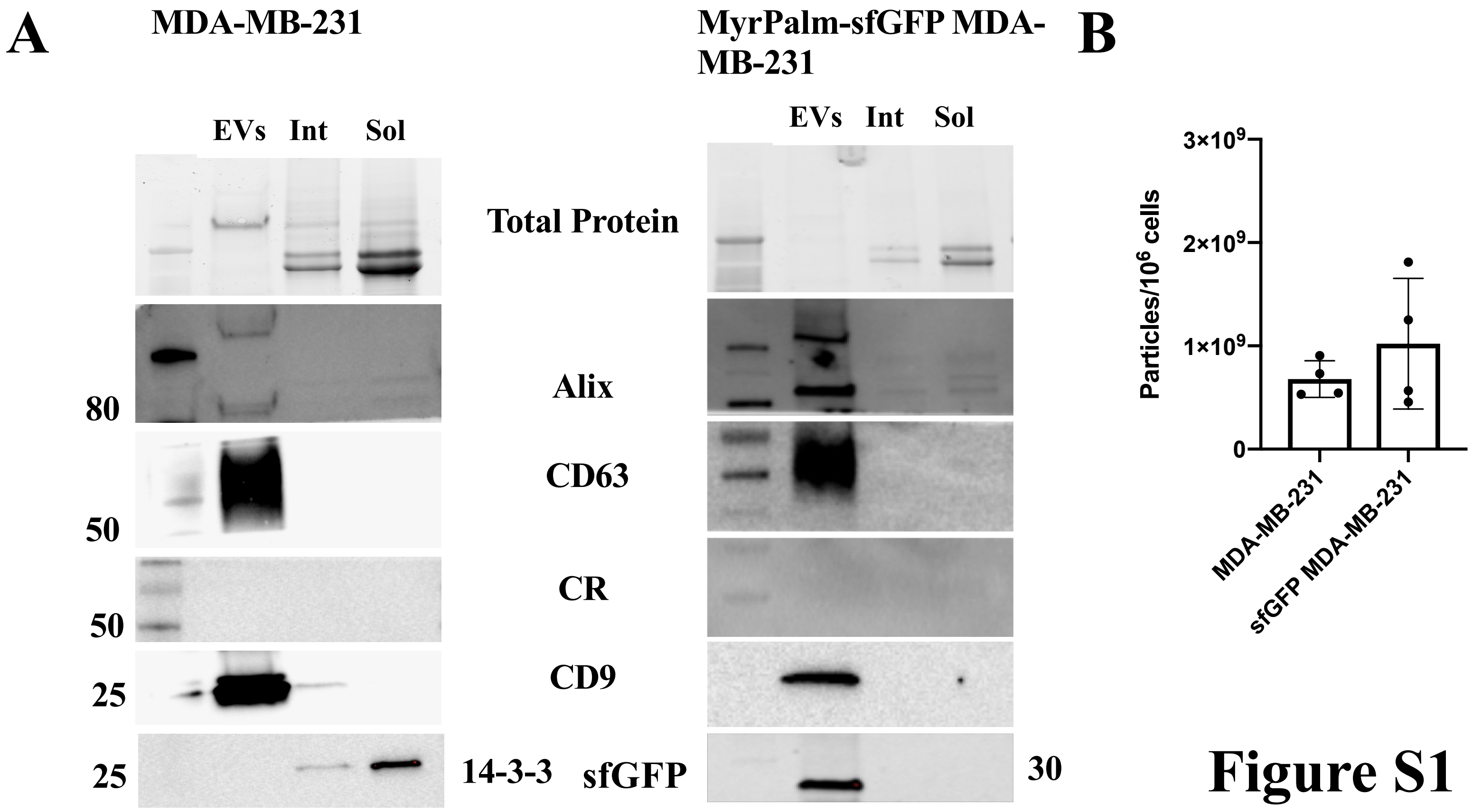
A
MDA-MB-231
MyrPalm-sfGFP MDA-MB-231
EVs
Int
Sol
Total Protein
Alix
CD63
CR
CD9
14-3-3
sfGFP
80
50
50
25
25
30
B
Particles/10^6 cells
3×10^9
2×10^9
1×10^9
0
MDA-MB-231
sfGFP MDA-MB-231

### Figure S1

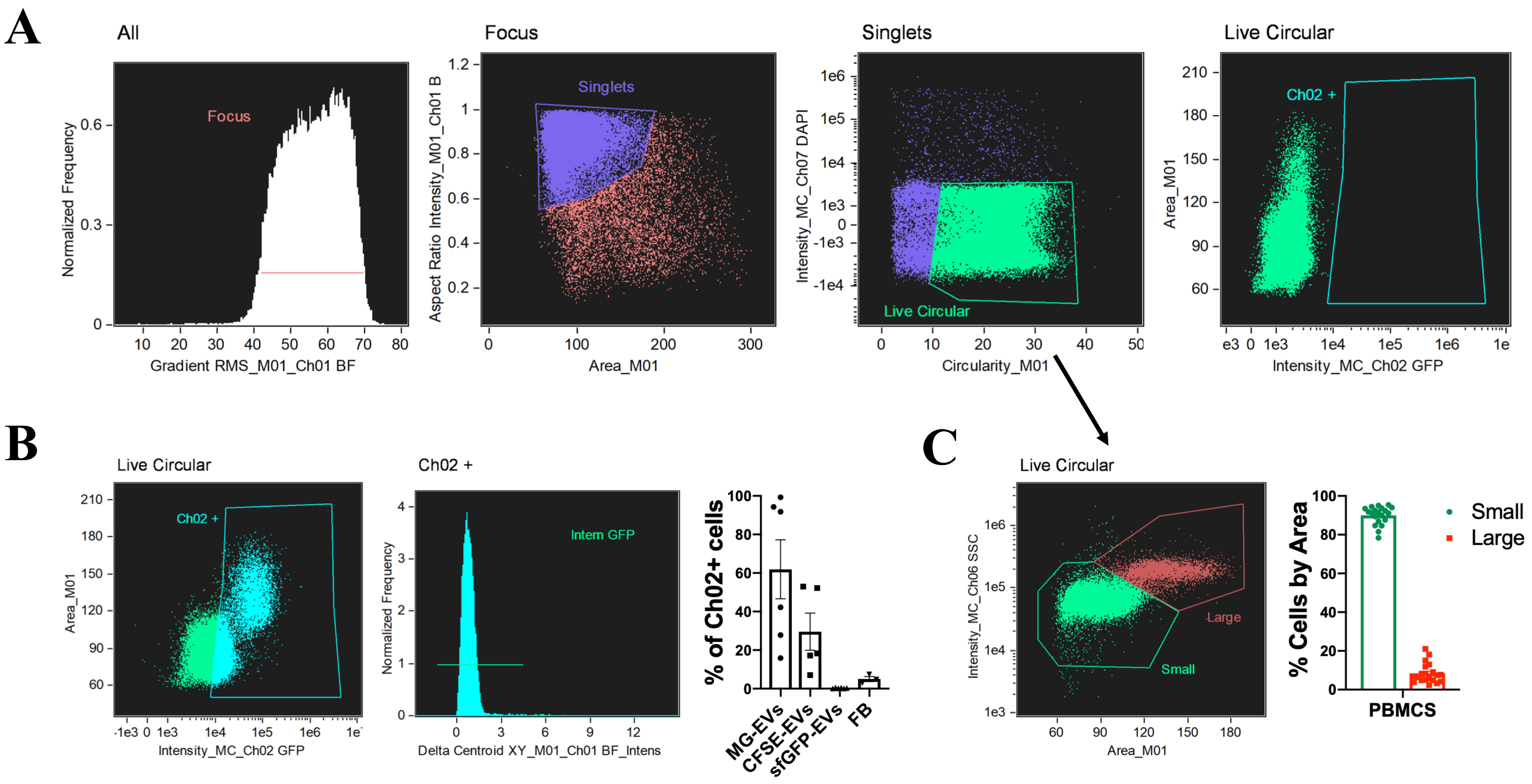
A
All
Focus
Normalized Frequency
Gradient RMS_M01_Ch01 BF
Focus
Singlets
Aspect Ratio Intensity_M01_Ch01 B
Area_M01
Singlets
Intensity_MC_Ch07 DAPI
Live Circular
Circularity_M01
Live Circular
Ch02 +
Area_M01
Intensity_MC_Ch02 GFP
B
Live Circular
Ch02 +
Area_M01
Intensity_MC_Ch02 GFP
Ch02 +
Intern GFP
Normalized Frequency
Delta Centroid XY_M01_Ch01 BF_Intens
% of Ch02+ cells
MG-EVs
CFSE-EVs
sfGFP-EVs
FB
C
Live Circular
Large
Small
Intensity_MC_Ch06 SSC
Area_M01
% Cells by Area
Small
Large
PBMCS

Figure S2

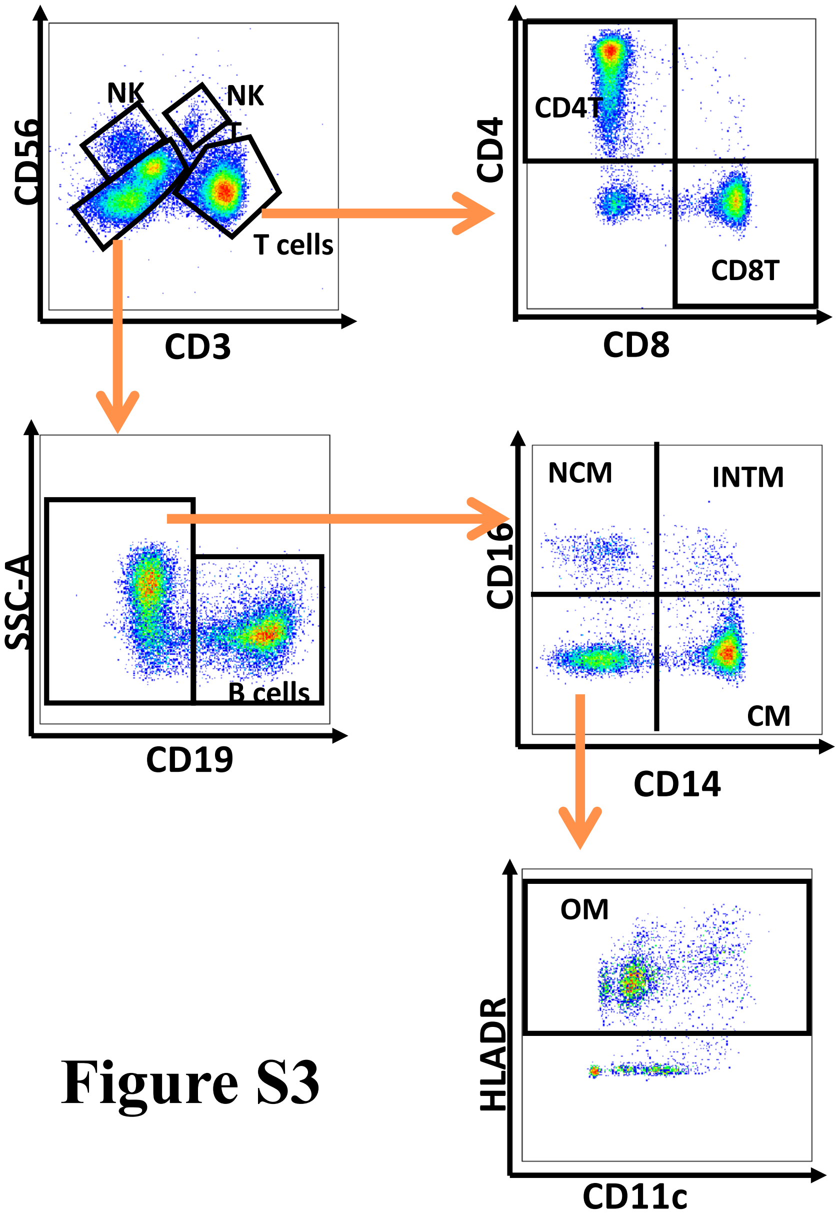
NK
NK
T
T cells
CD56
CD3
CD4T
CD8T
CD4
CD8
SSC-A
B cells
CD19
NCM
INTM
CM
CD16
CD14
OM
HLADR
CD11c

### Figure S3

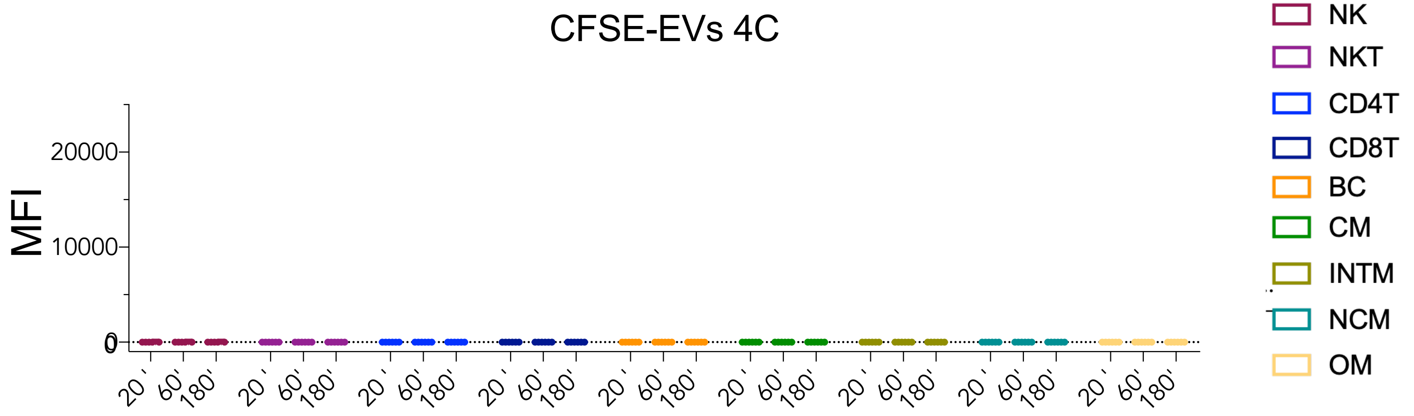
CFSE-EVs 4C
MFI
20000
10000
0
20'
60'
180'
NK
NKT
CD4T
CD8T
BC
CM
INTM
NCM
OM

**Figure S4**

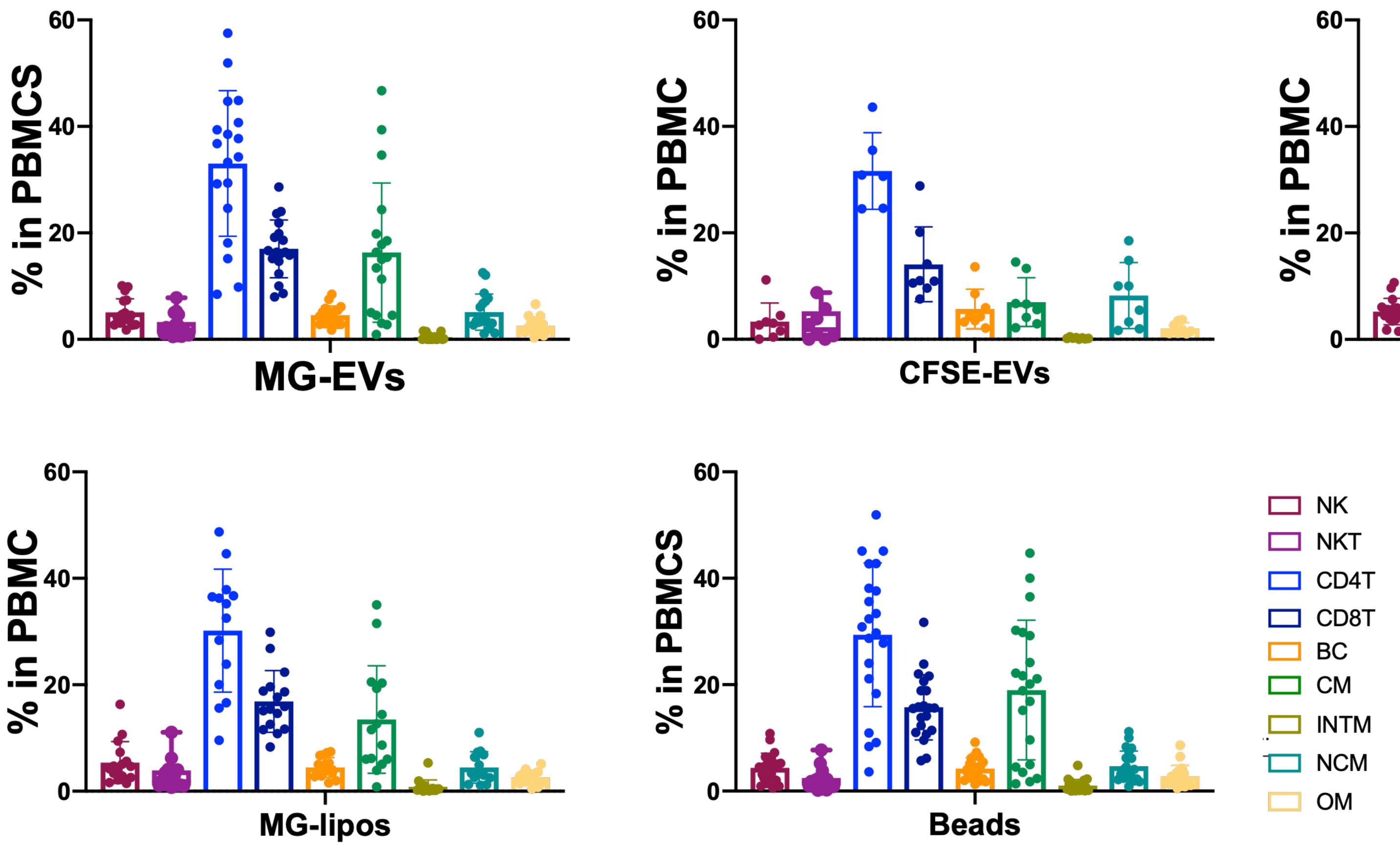
% in PBMCS
% in PBMC
MG-EVs
CFSE-EVs
sfGFP-EVs
MG-lipos
Beads
0
20
40
60
NK
NKT
CD4T
CD8T
BC
CM
INTM
NCM
OM

Figure S5
